# Helical grooves on the surfaces of microtrichia in European Hornets: Functional significance for antipodal relation between right and left hindwings

**DOI:** 10.1101/2024.02.25.582003

**Authors:** Sayako Inoué, Hisako Sato, Akihiko Yamagishi

**Affiliations:** Geodynamics Research Center (GRC), Ehime University, Matsuyama 790-8577, Japan; Faculty of Science, Ehime University, Matsuyama 790-8577, Japan; Faculty of Medicine, Toho University, Tokyo, 143-8540, Japan

## Abstract

The surface structures of microtrichia on the hindwings of *Verspa Crabro* (European hornets) were observed by scanning electron microscopy (SEM). Helical grooves were seen in the microtrichia on both the ventral and dorsal sides. Their helical orientation varied spatially across the wing surface but were the same in the ventral and dorsal surfaces. Notably the grooves wound antipodally between the left and right wings. The observed chirality relation might be related to the effective anti-wetting of hindwings. The results demonstrated the importance of microscopic chirality in understrading the functions of insect wings.

## Introduction

The surface morphology and topology of insect wings are important factors to control the functionality of insects. The three-dimensional micro-structures on the surface are responsible to the functionalities including but not limited to adhesion, locomotion, mechanical signal detection and hydrophobicity^1^.

Insect wings are usually covered by the dense arrays of microstructures such as bumps, scales, and hairs^2^. These structures have attracted much attention due to its application to bioinspired material developments^1,3^. The microtrichium is one of the hair-like structures commonly observed on the insect wings. The distribution and orientation of microtrichia on the wing surface are strongly related to the ability of insect wing to shed water away from the wing itself and from the body^4^. The hair-like structures on the insect wings often exhibit the hierarchical microstructure on the surface such as groove and chord^4,5^.

Similar hierarchical microstructures found in the insect legs are considered responsible for its hydrophobicity^6,7^. Byun et al. (2009) reported that the nano-scale grooves on the setae of *Tabanus chrysurus* help surface to repel water efficiently. They also stated that the Homoptera which has no hierarchal structure on its setae shows hydrophobicity due to its small size. The influences of hierarchical structure on the wing hairs, however, are still debatable. This situation is partly because of the very fine scale of a wing surface structure.

In this study, we found by scanning electron microscopy (SEM) observations that helical grooves cover the surfaces of microtrichia on the hindwings of *Verspa crabro flavofasciata*. By characterizing the morphological features of helical grooves, the complexity of surface microstructure was revealed with a focus on the functional significance. This study demonstrated the importance of microscale characterization of insect wing hierarchical structures particularly from the viewpoint of chirality.

## Samples and Methods

### Insect wings

The left and right hindwings of *Verspa crabro flavofasciata* Cameron 1903 (female) caught in Imabari, Ehime, Japan were observed in this study. The sample was endangered species donated by the Biodiversity Center, Ehime Prefectural Institute of Public Health and Environmental Science. This was a dead-body sample.

### Scanning electron microscopy

Both sides of wing surfaces were coated with gold using a JEOL JFC-1600 coater at the sputtering current of 20 mA for 60 seconds. The samples were observed by a field-emission scanning electron microscope (JEOL JSM-IT500HR) operated at 3kV.The obtained SEM images were analysed using Fiji (imageJ)^8^. The direction of helices was determined from the SEM image. The diameter and pitch of helices were measured from the hair on which the long axis appeared perpendicular to the viewing direction.

## Results and discussion

### Microtrichia on the ventral surfaces of hindwings

The SEM images of the ventral surface of the left and right hindwings are shown in Figures 1a and b, respectively. The regions of veins and membranes are visually distinguishable. The microtrichia cover the membrane part but sparingly exist on the surface of the vein. The microtrichia are grown towards the apical direction of the hindwing. The venation and orientation of microtrichia of the left and right hindwings appear symmetrical (Figures 1a and b). On the surface of each microtrichium, the helical groove is observed. The length of hair and orientation, the number of turns and the pitch of helices were determined on the SEM images (Tables 1 and 2).

**Figure 1.**
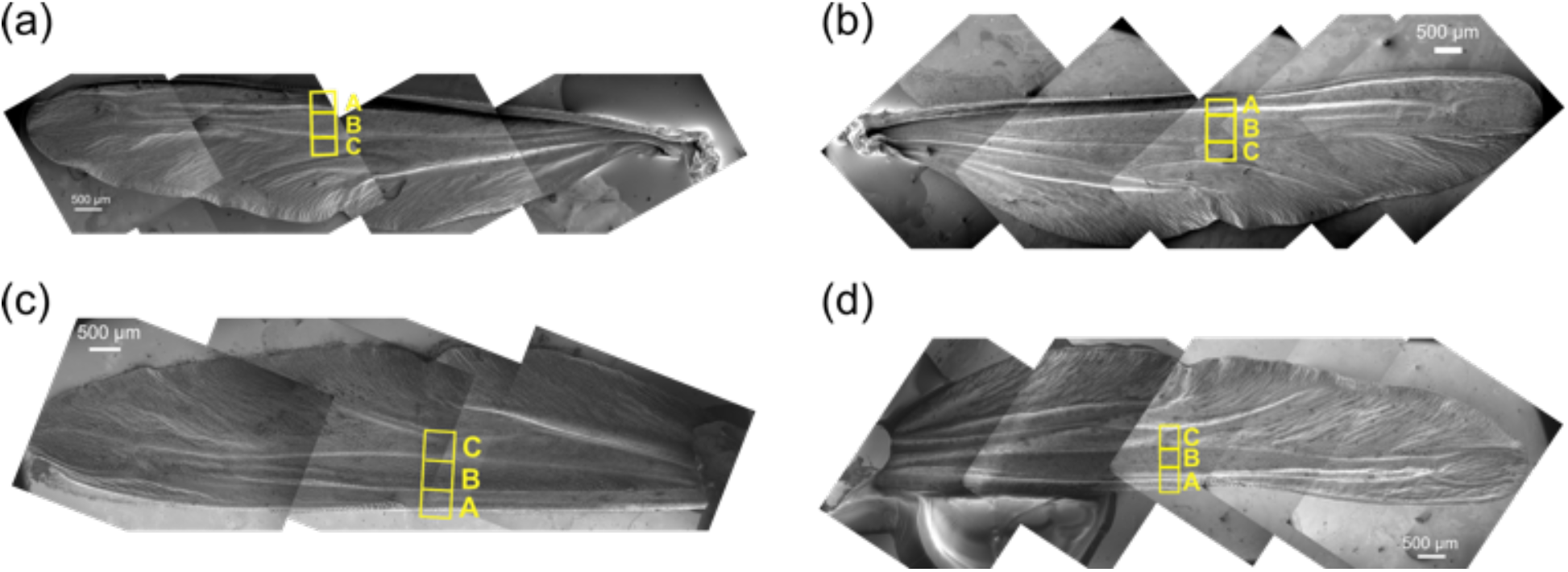
(a, b) Ventral surfaces of (a) left and (b)right hindwings. (c,d) Dorsal surfaces of (c) left and (d) right hindwings. The location of domain A, B and C was indicated in the figure.

**Table 1.** Average of length and longest diameter of microtrichia on ventral and dorsal surfaces of left and right hindwings. L: left wing, R: right wing, *n*: number of microtrichia analysed, σ : standard deviation, ^*^Only one hair was measured.

|  | Ventral |  |  |  |  |  | Dorsal |  |  |  |  |  |
| --- | --- | --- | --- | --- | --- | --- | --- | --- | --- | --- | --- | --- |
|  | Domain A |  | Domain B |  | Domain C |  | Domain A |  | Domain B |  | Domain C |  |
|  | L | R | L | R | L | R | L | R | L | R | L | R |
| *n* | 3 | 2 | 5 | 3 | 3 | 2 | 3 | 2 | 6 | 2 | – | 2 |
| Average Length<br>($\mu\text{m}$) | 86.5 | 95.4 | 95.6 | 80.8 | 93.0 | 90.3 | 102.9 | 87.5 | 58.6 | 55.2 | 48.2<br>* | 45.1 |
| $\sigma$ | 6.7 | 7.8 | 5.1 | 10.6 | 21.6 | 8.4 | 10.8 | 5.3 | 10.2 | 1.0 | – | 5.8 |
| Average longest<br>diameter ($\mu\text{m}$) | 14.3 | 14.4 | 17.2 | 14.7 | 18.6 | 18.7 | 24.6 | 22.8 | 17.4 | 8.4 | 7.3 * | 9.1 |
| $\sigma$ | 2.3 | 1.6 | 5.0 | 3.1 | 8.1 | 2.8 | 5.0 | 0.3 | 18.5 | 1.1 | – | 0.9 |

**Table 2.** Average of pitch of helical grooves observed on the microtrichia in figures 5 and 7. σ : standard deviation.

|  |  | Left wing | Right wing |
| --- | --- | --- | --- |
| Ventral surface | Pitch ($\mu\text{m}$) | 2.2 | 2.66 |
| | $\sigma$ | 0.54 | 0.44 |
| Dorsal surface | Pitch ($\mu\text{m}$) | 1.74 | 1.89 |
| | $\sigma$ | 0.41 | 0.33 |

Based on the orientation of helices, the observed area was divided into three domains: domain **A, B** and **C** (Figure 1). Domain **A** corresponds to the region in the cell nearest to the leading edge. Domains **B** and **C** are in the cells in the medial part of the wing. Domains **B** and **C** are divided by the longitudinal vein between two cells.

In domain **A** of the left hindwing, *right-handed* helical grooves were observed on the surfaces of microtrichia (Figure 2). Both right- and *left-handed* helices were observed in domains **B** and **C**. In these domains, however, the *left-handed* helices were more frequently observed. On the right hindwing, contrarily, *left-handed* helical grooves were observed in domain **A**. *Right-handed* helical groove was observed in domains **B** and **C** (Figure 3). Comparing the results between the left and right hindwings, it was deduced that the helical orientation of grooves was strictly antipodal in domain **A** and less strictly in domains **B** and **C**. In the present study, only chordwise distribution of helices was analysed. Still, the lack of consistency in domain **B** and **C** of left hindwing implies the complex distribution of helix orientation on the ventral surfaces.

**Figure 2.**
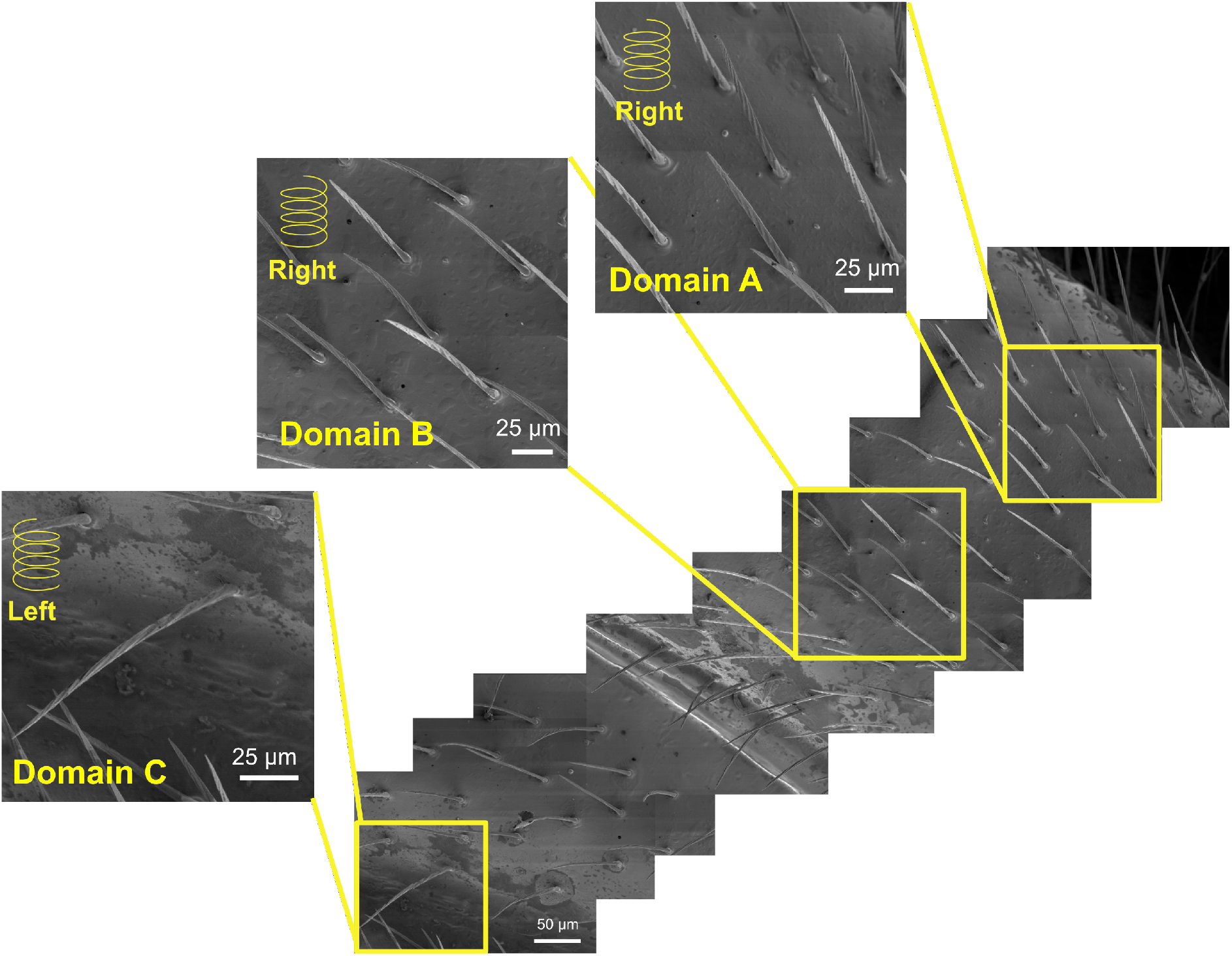
Microtrichia on the ventral surface of left hindwing. Domain A, B and C correspond to the position in Figure 1a. The orientation of helical groove is noted.

**Figure 3.**
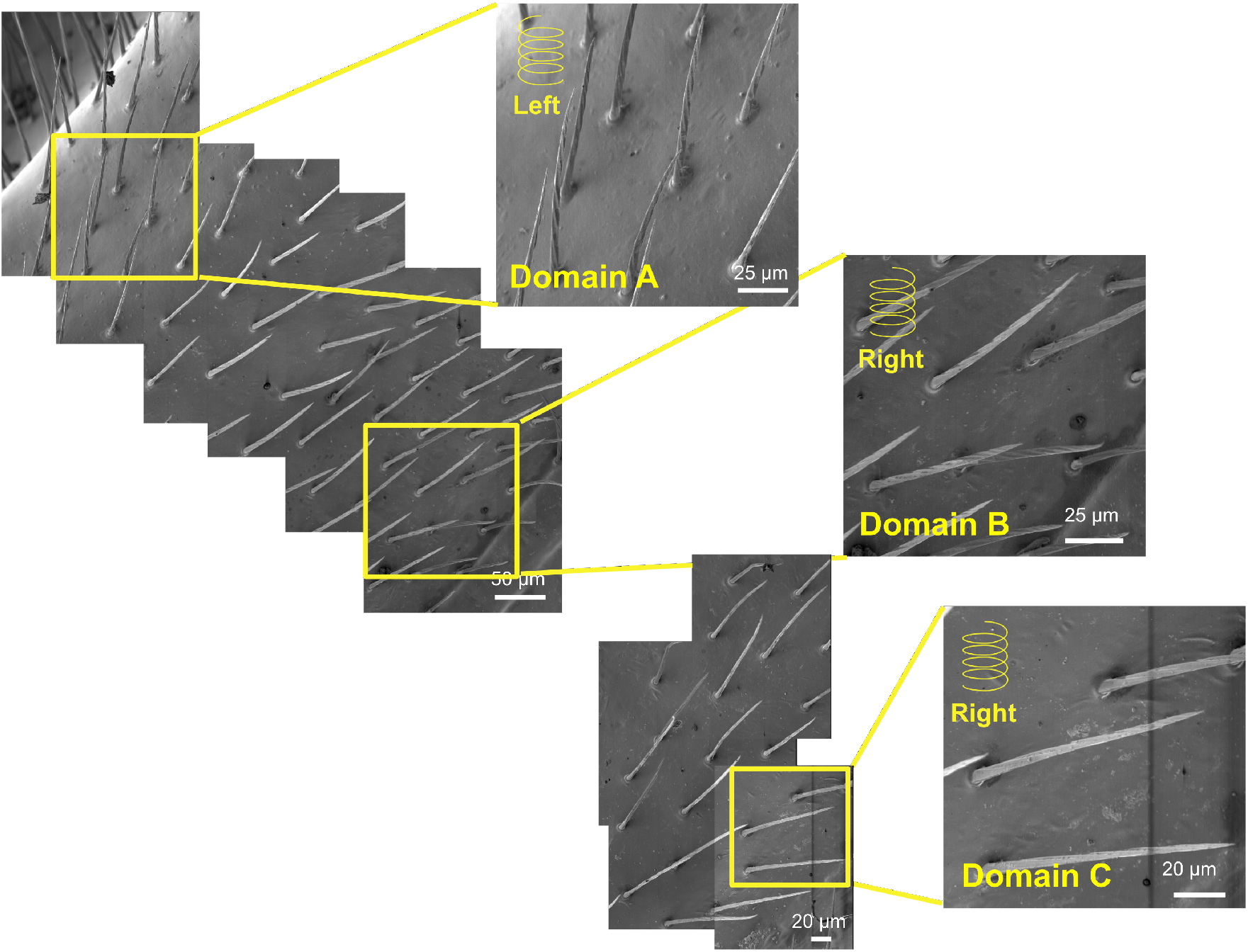
Microtrichia on the ventral surface of right hindwing. Domain A, B and C correspond to the position in Figure 1b. The orientation of helical groove is noted.

No correlation between the length of a hair and its helicity was observed. The number of turns in the helix was ranged between 5 and 11 for the left wing and 6 and 10 for the right wing, respectively (Figures 4a and b). The diameter of helices is not constant; the diameter is typically longest between 1/3 and 1/2 of the height from the root (Figures 5a-c). The trend in diameter indicates that the microtrichium has the slant cone shape or arcuate shape. It is difficult to estimate the degree of curvature from the SEM images because the viewing direction may not be perfectly perpendicular to the wing surfaces. Regardless of the length of hair, the longest diameter was constant (Figure 1S), implying the growth of hair was constrained crosswise. The pitch of helix is almost constant at 2 µm considering the accuracy of measurements on the SEM image (Figure 5d). The interpretation was supported by the trend that the number of helix turns got greater as the hair got longer (Figure 4).

**Figure 4.**
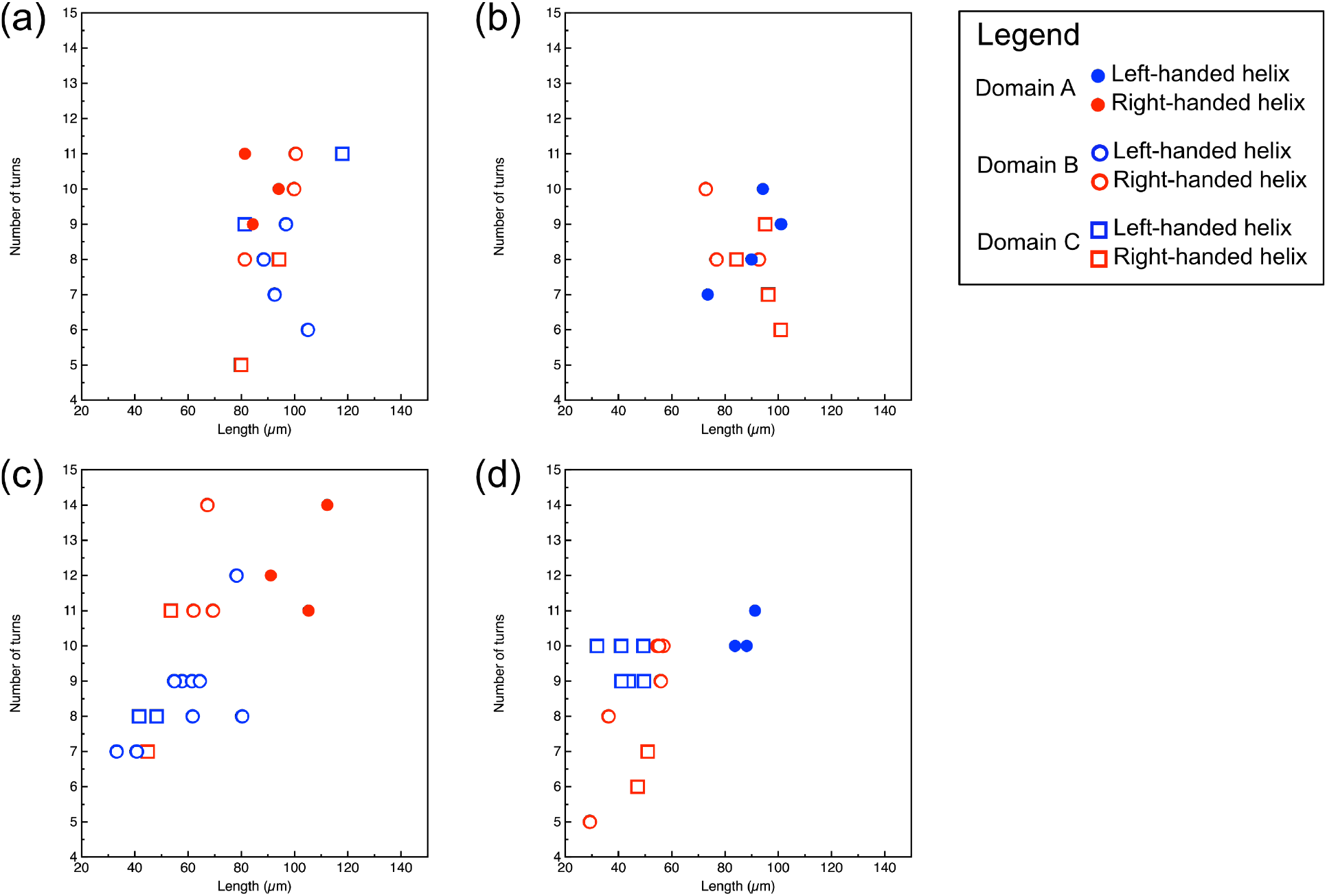
Plots of number of turns see in a microtrichium versus the length of hair on (a)left ventral surface, (b) right ventral surface, (c)left dorsal surface and (d) right dorsal surface.

**Figure 5.**
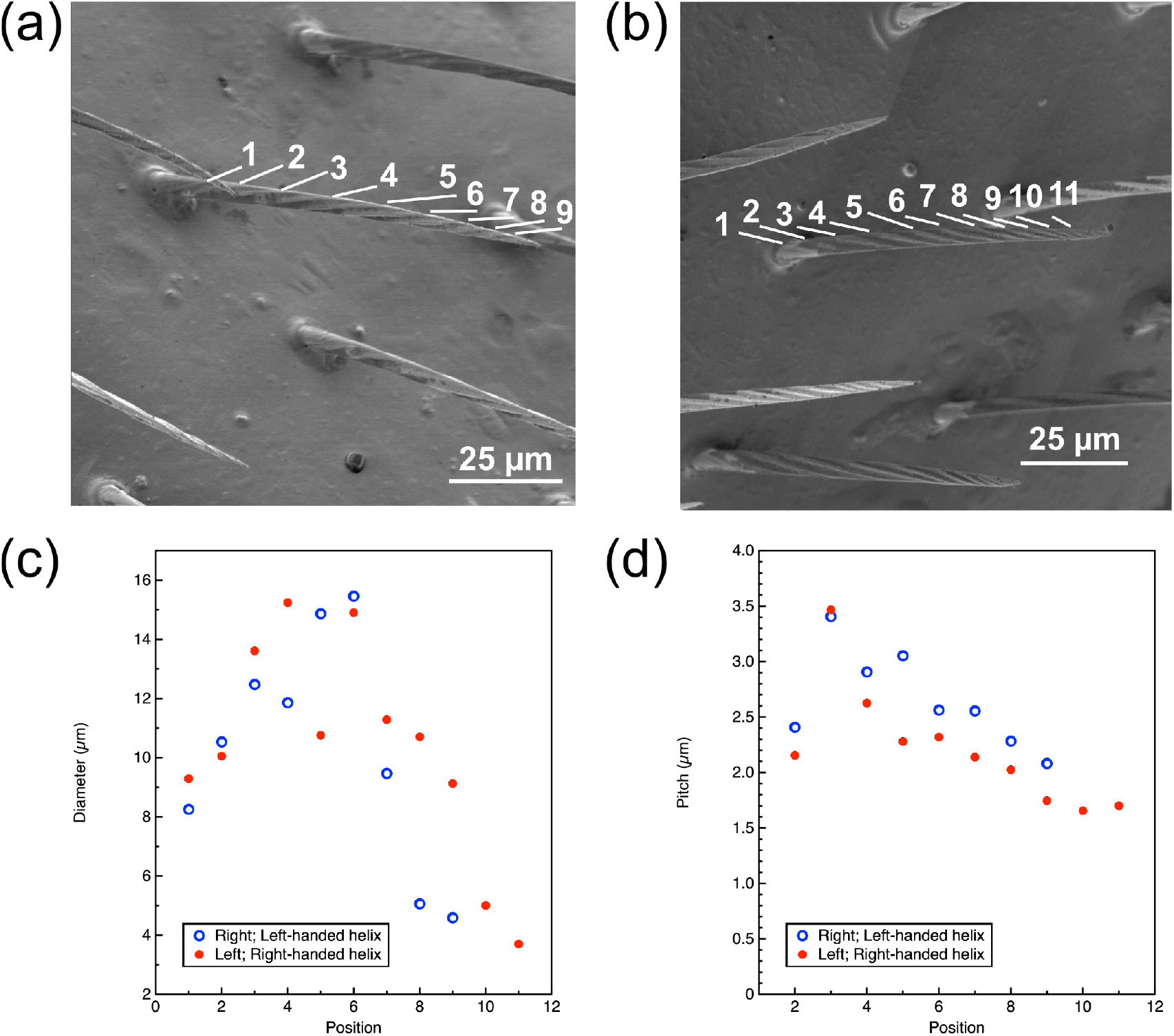
(a, b) Example of microtrichium on the ventral surface of left (a)and right (b) hindwings. Numbers show the positions of helix turn. (c) Plot of diameter along the turn vs position from the root of microtrichium. (d) Plot of pitch of a helix versus the position from the root. The pitch was measured as the distance between each helix turn. Position in figure c and d corresponds to the numbers on figure a, b.

**Figure 5.**
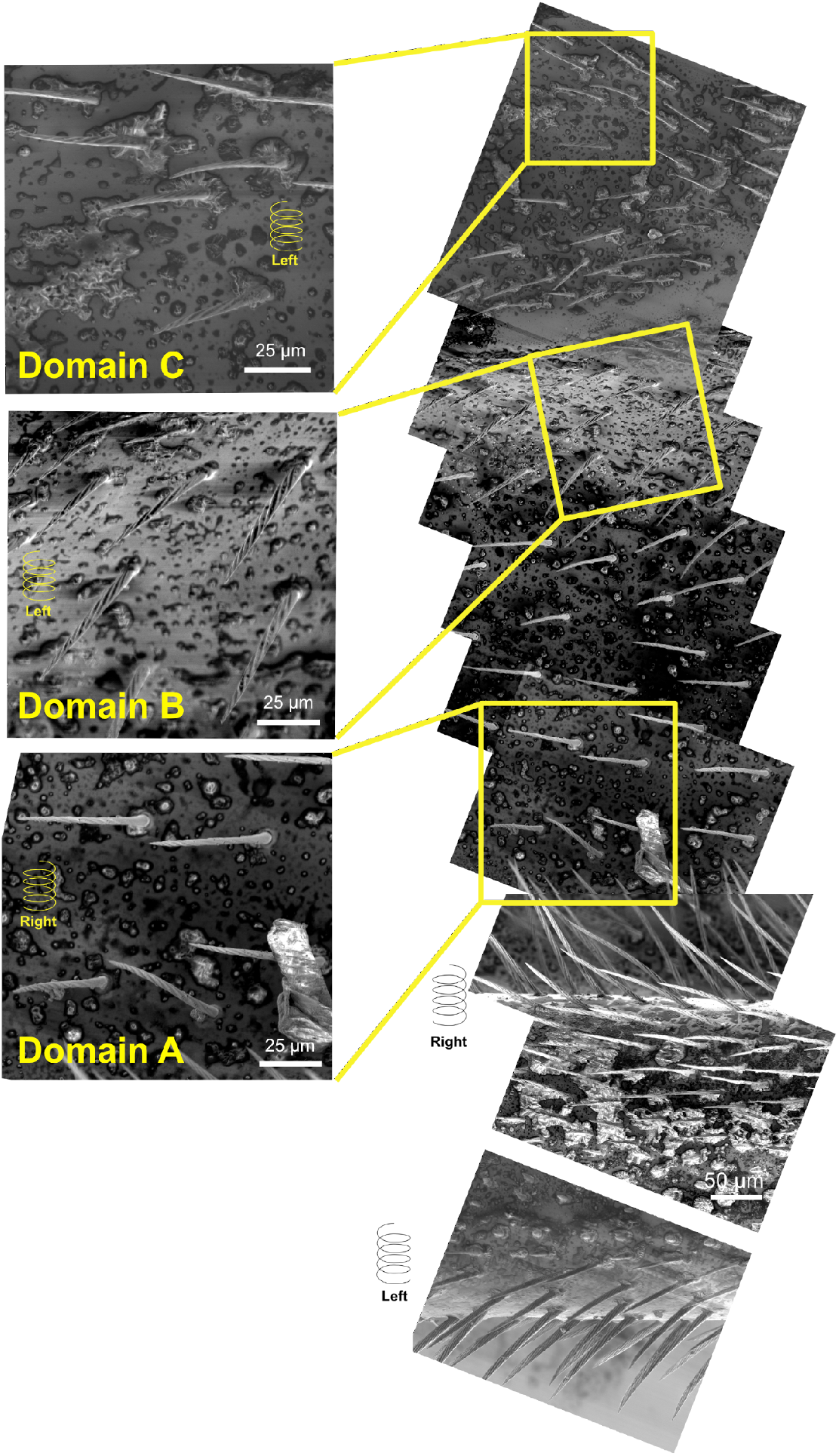
Microtrichia on the dorsal surface of left hindwing. Domain A, B and C correspond to the position in Figure 1c. The orientation of helical groove is noted.

### Microtrichia on the dorsal surfaces of hindwings

The SEM images of the dorsal surface of left and right hindwings are shown in Figures 1c and d, respectively. The microtrichia cover the entire surface of a wing. They were grown towards the apical direction of the hindwing. The helical groove is also observed on their surfaces (Figures 5 and 6). In contrast to the ventral surface, however, the distinction between vein and membrane regions is less clear. Nevertheless, the observed area was divided into three domains, domain **A, B** and **C**, by comparing the position with the ventral surface (Figures 1 c and d). Viewing from the dorsal side, the orientation of microtrichia of the left and right hindwings also appeared symmetrical (Figures 1c and d)^9^.

**Figure 6.**
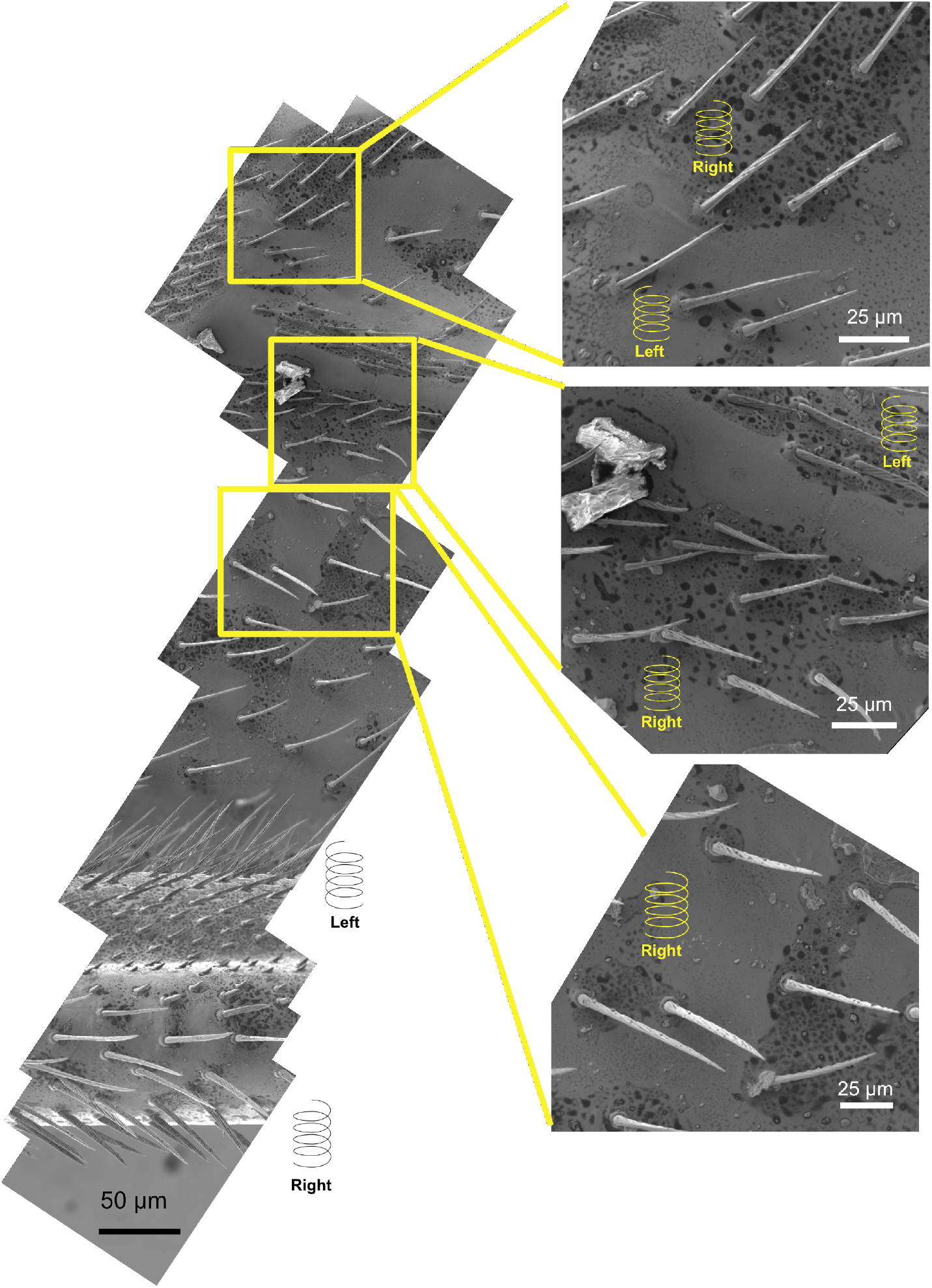
Microtrichia on the dorsal surface of right hindwing. Domain A, B and C correspond to the position in Figure 1d. The orientation of helical groove is noted.

On the left wing, the microtrichia have *right-handed* helical grooves in domain **A** (Figure 5). In the domains **B** and **C** of the left wing, both *left-handed* and *right-handed* helices were observed (Figure 5). The l*eft-handed* helices were more frequently observed in both domains. Contrarily the microtrichia in the domain **A** of the right wing have *left-handed* helical grooves (Figure 6). The *right-handed* helix was more frequently observed in the domain **B**. The occurrence of *left-handed* helix was more frequent in the domain **C**.

Because of the shape of wing, the hairs on the leading edge were able to observe from the dorsal side. The hairs on the edge have the *left-handed* helices in the left wing and *right-handed* helices in the right wing (Figures 5 and 6). In both cases, the helical orientation on the edge was opposite of that observed in domain **A**. The results of observation indicated that the helicity of grooves on the dorsal surfaces was also opposite between the right and left wings except for domain **C**.

On the dorsal surfaces of left and right wings, the average length of microtrichia in the domain **A**, *i*.*e*. near the leading edges of wings, was longer than the other domains (Table 1). The microtrichia in the domains **B** and **C** are shorter than those in the ventral surfaces (Figure 4). When compared between domains **B** and **C**, the length was similar. No clear correlation between length and helix orientation was observed (Figure 4). The number of turns in a helix was ranged between 7 and 14 for the left wing, and between 5 and 11 for the right wing. The number of turns increased as the hair got longer. The variance in the diameter of helices was also observed on the dorsal surface. In the case of both left and right wings, the diameter became longest around 1/3 and 1/2 of the height, suggesting the arcuate shape (Figure 7). The pitch of helix is almost constant at 1.8 µm. Positional relationships between microtrichia on the ventral and dorsal surfaces were undetermined as the cross-sectional view was not observed in this study. In the case of a male P. Heteroptera wing, the microtrichia grow at the same position on both sides of the wing surface ^4^.

**Figure 7.**
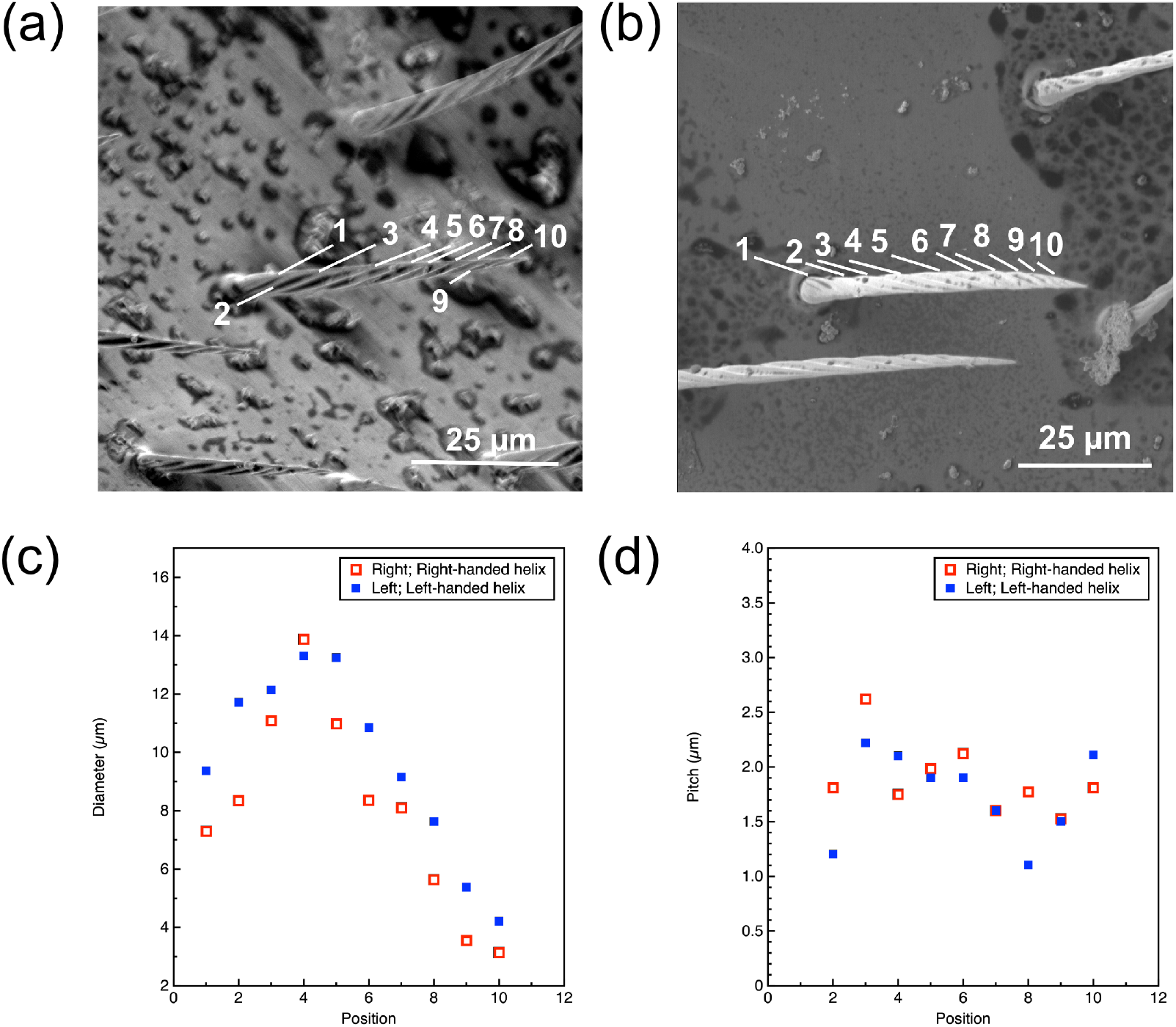
(a, b) Example of microtrichium on the dorsal surface of left (a)and right (b) hindwings. Numbers show the positions of helix turn.(c) Plot of diameter along the turn vs position from the root of mircrotrichium. (d) Plot of pitch of a helix versus the position from the root. The pitch was measured as the distance between each helix turn. Position in figure c and d corresponds to the numbers on figure a, b.

### Implications of helical groove on the microtrichia

The results on the helicity of a groove in each domain are summarized in Figure 8 and Table S1. The helical grooves on the ventral and dorsal surfaces of hindwings showed the same orientation except for the domain **C** (Table 2). However, the difference in the domain **C** is likely to arise from the complex spatial distribution of helix orientation. Therefore, we concluded that the orientations of helices are the same on ventral and dorsal surfaces. The curvature of microtrichium seems to be related to the helix pattern on the surface. If the helical pattern on the surface indicates the growth direction of the microtrichium, it is reasonably assumed that the growth direction of hair is controlled by the orientation of helix.

**Figure 8.**
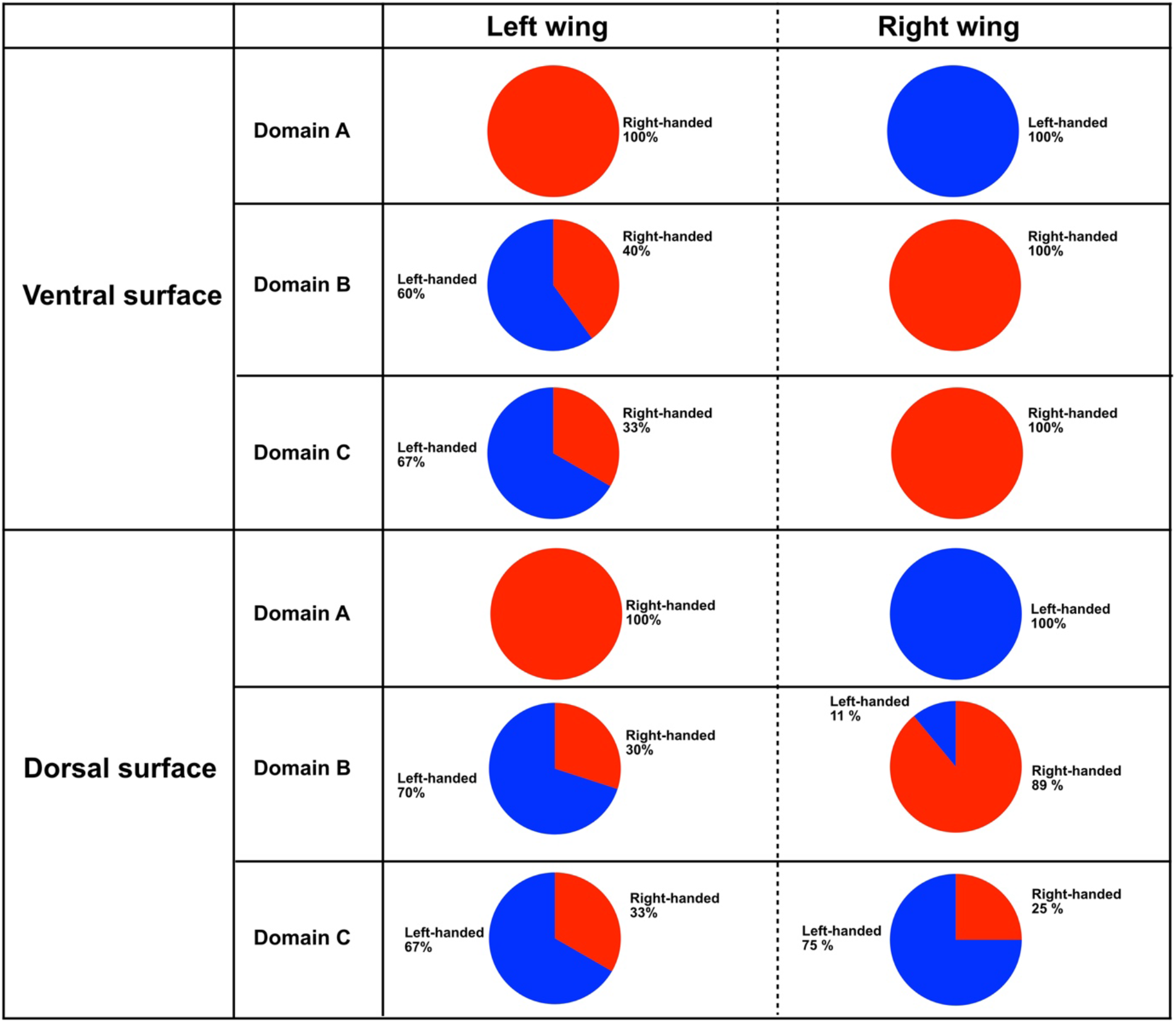
Summary of helicity of grooves. Pie charts indicate the frequency of helix orientation observed.

The chirality in a helical groove may be responsible for satisfying the symmetrical spatial distribution of microtrichia on left and right wings. The distribution patterns of microtrichia are related to their functions. Firstly, they could provide stable interlocking structures. In most Hymenoptera species, the forewing and hindwing are coupled to each other during flight by hooks at edges and act as one wing^10–12^. The hooks called hamuli on the anterior margin of the dorsal side of hindwing are mainly responsible for the coupling of two wings ^13,14^. In our study, the length of microtrichia on the dorsal surface is longer in the area nearest to the anterior margin than the other part of wing. This difference may reflect their contribution to the interlocking structure.

The distribution patterns of microtrichia also have a significant impact on the anti-wetting property of a wing ^4–6^. The microtrichia are oriented in response to the topology of the wing surface to shed water away from the body and wing^4^. The existence of helical groove itself on the microtrichium surface would also contribute to the functionality of the wing. The roughness of microtrichium surface should help surfaces repel water more efficiently as the surface grooves create the air pockets at the interface between water and microtrichium^15^. The helical groove would allow water to move away from the microtrichium. Together with the orientation of microtrichia direction, the helical grooves may reinforce the anti-wetting function of microtrichia.

Finally, it is emphasized that the present study is the first report on the helical pattern on the insect wing microtrichia. The helical microstructure would create micro vortex on the insect wing surface, which may have impact on the aerodynamic properties. A preference of one chiral form to another has often been observed in the building blocks of living things^16^. It is no surprise if the existence of chirality relation in the wing surface microstructure may be much more common in nature than we presently assumed. Detailed characterization of micro-scale structures on the insect wings will help in understanding their functionality.

## Conclusion

We reported the helical pattern on the surface of microtrichia on the hindwings of *Verspa Crabro* (female). The analyses of helix orientation on the ventral and dorsal surfaces of left and right hindwings revealed that they are in a chiral relationship. On the same side of the wing, the trend in helix orientations differs depending on the area on the wing. The microtrichia near the leading edge of hindwings have the helix orientation opposite from the inner part of the wing. Although the area analysed in this study was limited, the complex distribution of helix orientation was suggested. The helical pattern may be related to the orientation of microtrichium and wetting behaviour of wing.

## Acknowledgements

The authors are grateful to Mr. Katsushi Narimatsu and Mr. Yusuke Hara (Biodiversity Center, Ehime Prefectural Institute of Public Health and Environmental Science) for donating *Vespa crabro flavofasciata*. This study was supported by the Japan Society for the Promotion of Science (JSPS, KAKENHI (grant number: JP22H02033). This study was supported by the Joint Usage/Research Center of Premier Research Institute for Ultrahigh-pressure Sciences (PRIUS, Project No. 2023-D08) at Ehime University, Japan.

