## Supplemental information for "Helical grooves on the surfaces of microtrichia in European Hornets: Functional significance for antipodal relation between right and left hindwings"

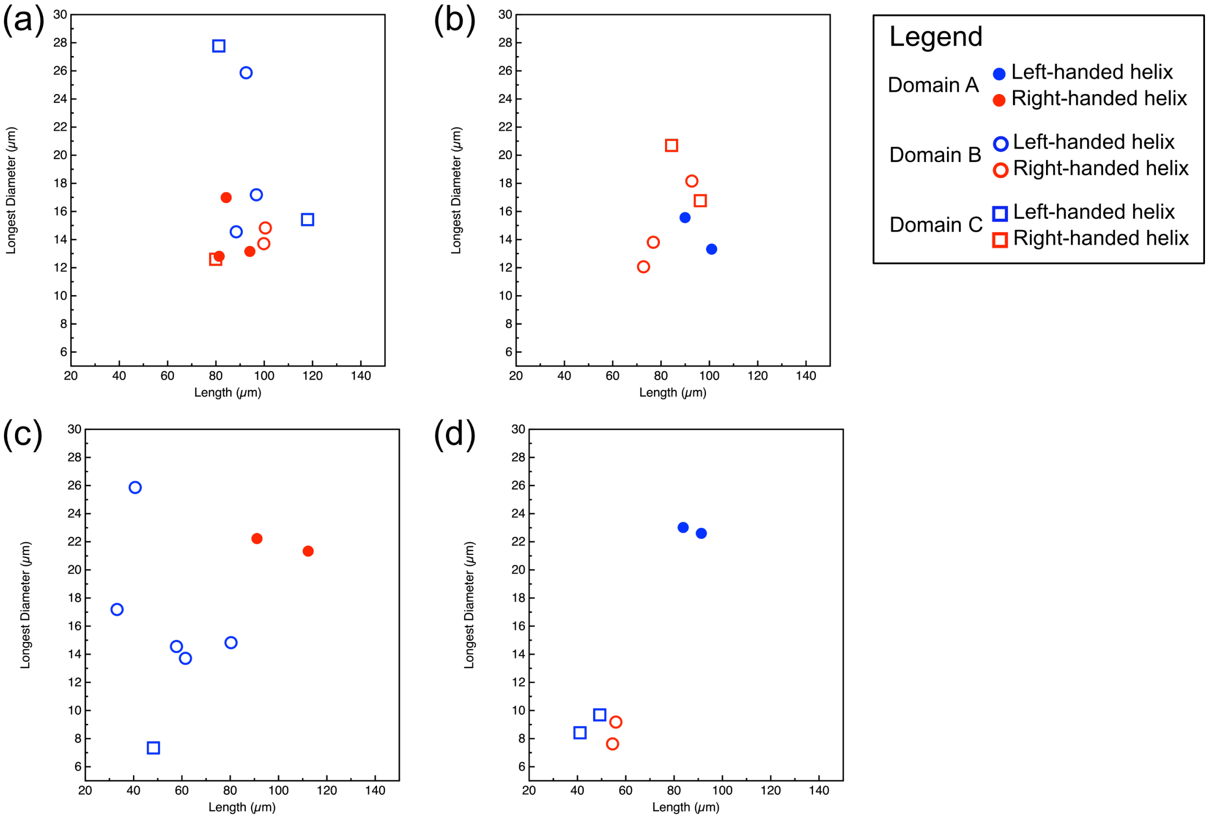


**Figure S1. The plots of longest diameter of helical groove versus length of hair. (a) Ventral surface of left hindwing. (b) Vental surface of right hindwing. (c) Dorsal surface of left hindwing. (d) Dorsal surface right hindwing.**

| | Ventral | | | | | | Dorsal | | | | | |
|---|---|---|---|---|---|---|---|---|---|---|---|---|
| | Domain A | | Domain B | | Domain C | | Domain A | | Domain B | | Domain C | |
| | L | R | L | R | L | R | L | R | L | R | L | R |
| Left-handed | 0 | 5 | 4 | 5 | 4 | 5 | 0 | 3 | 9 | 1 | 4 | 6 |
| Right-handed | 3 | 0 | 3 | 0 | 2 | 0 | 3 | 0 | 3 | 8 | 2 | 2 |

**Table S1. The orientation of helices observed the ventral and dorsal surfaces of left and right hindwings. Numbers on the table indicate the number of microtrichia with left-or right-handed helix observed. L: left wing, R: right wing.**
